# Correlative MIMS-EM imaging reveals metabolic turnover from organelle to organismal scales in *C. elegans* during dietary restriction

**DOI:** 10.64898/2026.08.20.746106

**Authors:** Alessandra Norris, Christopher Acree, Li Peng, Rafael Arrojo e Drigo, Kristopher Burkewitz

## Abstract

Metabolism is spatially compartmentalized across scales, from distinct tissues to cells and organelles. However, most approaches for studying metabolic activity obscure spatial organization and intra-compartment heterogeneity within bulk biochemical measurements. On the other hand, multi-isotope mass spectrometry coupled with scanning electron microscopy (MIMS-EM) maps the fates of labeled nutrients *in situ* at nanometer-scale resolution, preserving ultrastructural context. Here we adapt MIMS-EM for *Caenorhabditis elegans*, where the compact metazoan body plan uniquely enables visualization of virtually all tissue types and their resident organelles within a single cross-sectional image. Using pulse-chase labeling of dietary carbon and nitrogen, we apply this approach to understanding the metabolic program induced in early stages of dietary restriction (DR). While DR is widely proposed to enhance organismal healthspan by enhancing broadscale turnover, proteomic studies have suggested more nuanced models. MIMS-EM across intact animals reveals that DR induces non-uniform effects between tissues and carbon/nitrogen resources, accelerating carbon turnover in the muscle and hypodermis, but not intestine. At the organelle scale, MIMS-EM revealed heterogeneity within mitochondrial networks that was independent of diet and stable over time. Spatial analysis of isotope signatures within intestinal mitochondrial networks also indicated greater similarity between neighboring mitochondria than distal mitochondria, supporting models of local mitochondrial mixing. Collectively, these results reveal that DR induces compartment- and resource-specific remodeling strategies across an intact animal while establishing *C. elegans* MIMS-EM as a powerful platform for multi-scale, integrative models of nutrient handling.

## Introduction

Effective distribution of nutrient resources is especially complex in metazoans, where metabolism is partitioned across a multi-scale array of tissues, cells, and organelles. Each of these compartments is highly specialized and interdependent on one another, necessitating continual coordination of their respective metabolic activities as physiological demands change. Efficient metabolic coordination across these scales gives rise to metabolic plasticity and resilience, whereas failures in this coordination contribute to diverse chronic and age-dependent diseases^1,2^. Understanding how nutrients are differentially allocated between organelles within a cell, between cells within a tissue, and between tissues within an animal is thus centrally important to our understanding of health and physiology. However, current models for understanding how nutrient allocation is integrated in an animal are limited by the inherent technical and analytical challenges of capturing and resolving data between so many compartments.

The challenge of understanding nutrient allocation strategies compounds at the smallest scales. Organelles are highly dynamic and respond to fluctuations in the environment by changing in size, shape, abundance, composition and spatial distribution in order to adapt and optimize function. Increasingly, we understand that through this repositioning of organelles, cells can effectively re-wire inter-organelle contact sites, signaling and metabolic flux^3,4^. These structural transitions are achieved in part through coordinated degradation and synthesis of organelle membrane and protein components. Consequently, nutrients entering the cell are not only allocated spatially between organelles to support the reactions of energy metabolism housed within them, but also for the biosynthetic and turnover pathways utilized for expanding and remodeling the organelles themselves.

Understanding the principles guiding how metazoans invest and turn over their nutrient resources requires methods capable of tracking nutrient fates while preserving the spatial organization of these compartments. While metabolomic and conventional isotope-tracing approaches have transformed our understanding of metabolic regulation, these methods rely on bulk measurements and fractionation strategies that obscure spatial organization and heterogeneity between individual cells and organelles. Multi-isotope mass spectrometry (MIMS)-based imaging overcomes these limitations by providing nanometer-scale resolution of stable isotope labeled-nutrients within intact specimens^5,6^. MIMS imaging can also be coupled with correlative scanning electron microscopy (EM), providing the ultrastructural background upon which isotope abundance can be quantitatively localized down to sub-organelle resolution^6^. While these strengths have been realized in isolated mammalian tissues^5,7^, the compact body plan of the *C. elegans* model provides an untapped opportunity to extend this framework across an entire metazoan. Because virtually all major tissue types of the animal are present within a single 50-100 m cross-section, a single, high-resolution MIMS-EM image enables visualization of stable isotopes not only across all major cell types of the animal, but also all the organelles within those cells. Combined with the rapid genetics, compressed lifespan, and conserved nutrient-responsive longevity interventions, this *C. elegans* MIMS-EM platform can theoretically provide empirical data defining nutrient allocation and turnover strategies from the organelle to the organismal scale across development and aging.

Dietary restriction (DR)-based interventions provide a compelling context in which to apply this multi-scale approach. DR confers dramatic extensions to healthspan and lifespan across many species^8–11^. The strength and consistency of this observation underscores a long-held hypothesis in aging biology that DR triggers an evolutionarily ancient metabolic reprogramming strategy, optimized for survival and resilience when food is scarce^12,13^. Over the past two decades, transcriptomic, proteomic and metabolomic studies have helped to define many of the molecular pathways contributing to the life-extending effects of DR, including evolutionarily conserved changes in nutrient-signaling pathways^8–11^ and remodeling of cellular organelles^14–18^. While the catalog of nutritional and molecular mediators of DR is well established, however, clear models describing how nutrient resources are differentially utilized across scales during DR have yet to emerge.

The challenge of defining such longevity-associated metabolic programs is compounded by multiple factors, including the complex, multi-scale array of compartments mentioned above, as well as the diversity of DR interventions, as nutrient composition and feeding timing can influence DR efficacy^19–21^. Nevertheless, several broad features of the DR response have emerged. Physiologic and multi-omic analyses have shown that metabolic rewiring occurs during DR in mammalian models, including changes in amino acid and lipid metabolic pathways^22,23^. At the cellular level, longevity is also consistently associated with reduced protein synthesis and increased autophagy^23–26^. In particular, this inverse relationship between biosynthetic and degradative processes has led to the paradigm that enhanced turnover and cellular recycling is a foundation of healthy aging^27,28^. However, tissue-focused studies of proteome turnover have countered this view of DR, showing that chronic DR is associated with more complex macromolecular alterations, including an unexpected reduction in turnover of many proteins in liver and muscle^29–31^. These findings suggest that rather than acting through a non-specific bulk recycling process, DR may selectively remodel particular molecular and organellar components while stabilizing others. These competing models have not been reconciled, and understanding how turnover processes are distributed spatially across tissues and subcellular compartments remains an important unresolved question.

Here we develop and apply a MIMS-EM-based, spatial pulse-chase approach for *C. elegans* and begin mapping how nutrient interventions remodel metabolic turnover across scales. We develop a sample processing pipeline to prepare the animals for MIMS-EM imaging, including a protocol for controlling ^13^C and ^15^N stable-isotope labeling of the *C. elegans* bacterial diet. From this protocol we demonstrate robust label incorporation and MIMS-EM resolutions corresponding generally with the practical maximum (∼50-60 nm for MIMS and 5 nm for SEM). Applying MIMS-EM to the context of DR, we identify tissue-specific differences in turnover kinetics across major somatic tissues. We find that DR accelerates carbon turnover in muscle and hypodermis, but not intestine, and that tissue-level nitrogen kinetics are partly uncoupled from carbon. We examine mitochondrial networks in the same cells, uncovering generally higher turnover rates relative to the host tissues, but similar turnover-accelerating effects of DR. Finally, we examine isotope distributions within the mitochondrial network of individual intestinal cells, revealing heterogeneity at single-organelle resolution. Spatial analysis of these mitochondrial isotope signatures provides support that metabolic variance within the network is buffered by localized mitochondrial mixing (e.g., fission/fusion), which equilibrates across the cell over time. Together, these results provide the basis for understanding how DR reprograms nutrient investment and turnover across tissues and organelles, while establishing MIMS-EM in *C. elegans* as a platform for multi-scale analysis of metabolic remodeling.

## Results

### Development of an efficient stable-isotope labeling strategy *C. elegans* labelling *in vivo*

In tissue culture and mammalian models where MIMS-EM has previously been applied, stable isotopes are typically incorporated through cell media supplementation, intravenous infusion, or feeding of defined labeled substrates, such as ^15^N-thymidine or [U-^13^C_6_]glucose^6,32^. However, the small size of *C. elegans* precludes infusion-like approaches, and the low permeability of the worm cuticle limits nutrient uptake by absorption from the media. To implement MIMS-EM in *C. elegans*, we thus focused on establishing a feeding-based protocol to introduce stable isotopes *in vivo*.

*C. elegans* feed on *E. coli* bacteria, which introduces an additional and dynamic biological system between labeled tracers and animal incorporation. To avoid the potential for asymmetric incorporation of ^13^C and ^15^N tracers among distinct macromolecular pools (e.g., lipids vs. carbohydrates and amino acids), which could influence downstream interpretation, we developed a protocol to saturate *E. coli* cells with ^13^C and ^15^N isotopes across all bacterial metabolites (**Figure S1A**). To accomplish this, we first optimized growth in a defined minimal media (MM), where[U-^13^C_6_] glucose and ^15^NH_4_Cl are added to M9 buffer to provide the only carbon and nitrogen sources. Notably, we initially found that growing OP50 in minimal media results in a substantial reduction in bacterial growth compared to LB media, which could potentially complicate downstream processes for preparing *C. elegans* diet restriction paradigms^33^ (**Figure S1B**). We addressed this problem by avoiding common autoclave-based sterilization step of the M9 buffer, and instead relying upon filtration, which increased bacterial yield (**Figure S1B**). As minimal media is nutritionally limited, we also evaluated a commercially available labeled complete media (CM), where all algae biomass (including metabolites) are saturated with ^13^C/^15^N isotopes. We found that this complete media improves bacterial growth compared to the defined MM (**Figure S1C**), highlighting its utility especially when large amounts of labeled bacteria are needed. We next evaluated whether feeding worms with bacteria grown in ^13^C/^15^N-saturated media causes gross physiological effects (**Figure S1D**). We allowed synchronized eggs to develop on CM, MM or heavy MM and found no differences in worm growth under these condtiions (**Figure S1E**). This indicates that worm feeding using isotopically labelled bacteria grown in CM and MM does not interfere with worm growth. Since no differences were observed when using labelled bacteria grown using CM or MM, we utilized the cost-effective MM for our downstream studies.

### Processing *C. elegans* samples for MIMS-EM

We next aimed to test the effectiveness of this labeling strategy on *C. elegans* and determine the practicality of imaging these samples with MIMS-EM. Synchronized eggs were placed on labeled lawns and allowed to develop until day 1 of adulthood before MIMS-EM processing (**Figure 1A**). In this experimental paradigm, embryonically derived biomass is unlabeled while post-embryonic contributions to organism biomass are expected to be composed of diet-derived ^13^C and ^15^N. As worms undergo an estimated ∼60-100-fold expansion between embryo and adult stages^34^, the theoretical proportion of embryonically derived biomass is thus limited to <2% overall.

**Figure 1.**
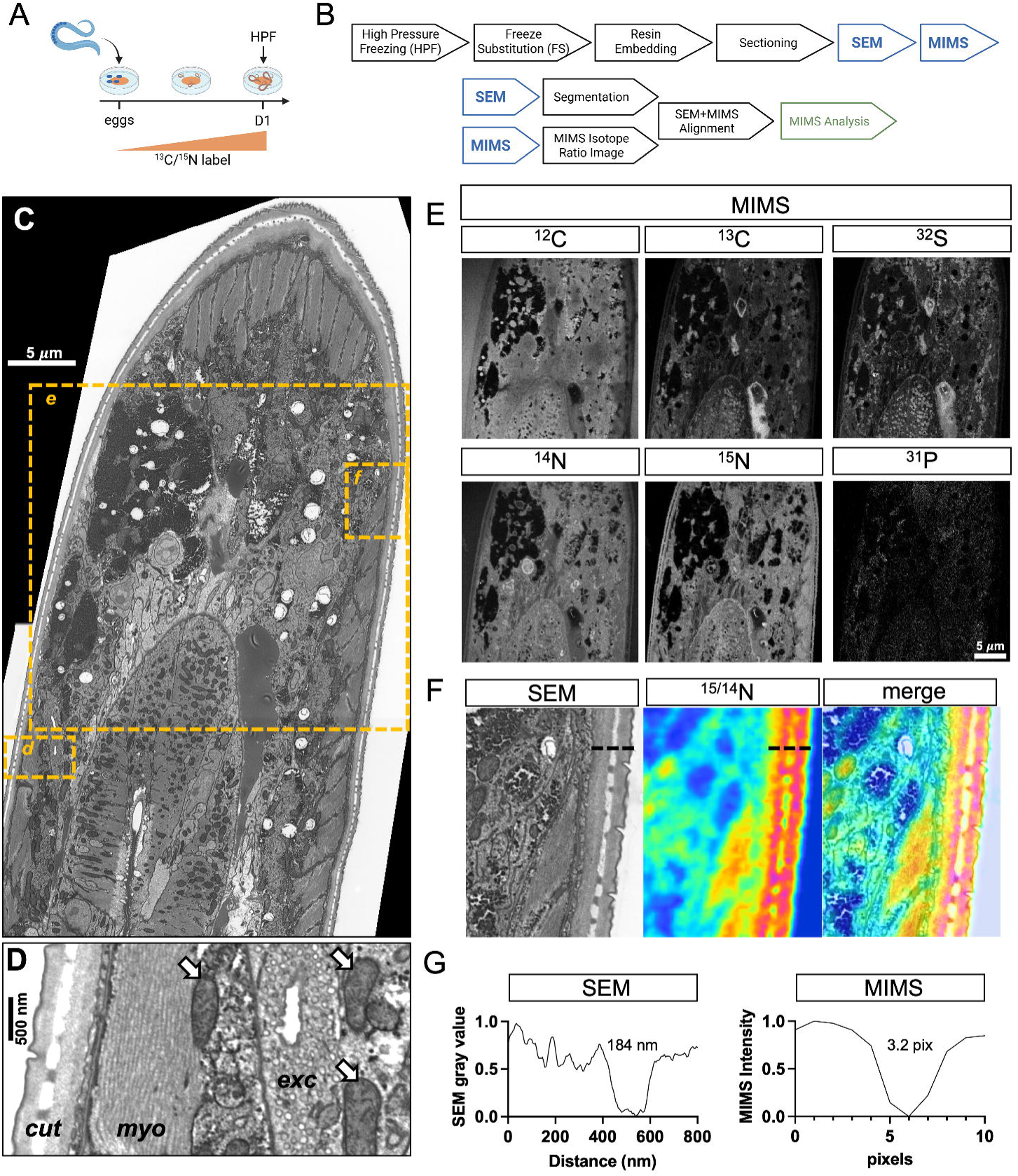
A correlative MIMS-EM workflow for *C. elegans*. (A) Labeling scheme. Synchronized eggs were placed on ¹³C/¹ □ N-labeled bacterial lawns and grown to the first day of adulthood (D1) prior to high-pressure freezing (HPF). (B) Sample processing and image analysis pipeline. Sections were imaged sequentially by SEM and MIMS, with segmentation performed on SEM images and isotope ratios extracted from registered MIMS images. (C) SEM cross-section through the anterior region of a D1 animal. Boxed regions are shown in (D) and (F). Scale bar, 5 µm. (D) Higher-magnification SEM of the region boxed in (C), showing the cuticle (cut), muscle myofilaments (myo), and excretory cell (exc). Arrows indicate mitochondria. Scale bar, 500 nm. (E) Secondary ion images of the region boxed in (C) for the indicated isotopes, acquired in parallel from a single MIMS field of view. Scale bar, 5 µm. (F) SEM, ^15/14^N ratio, and merged images of the cuticle and adjacent hypodermis. Dashed lines indicate the position of the line profiles shown in (G). (G) Intensity profiles across the cuticle from the SEM (left) and MIMS (right) images shown in (F), used to calibrate MIMS pixel size. Full width at half maximum was 184 nm in the SEM image and 3.2 pixels in the MIMS image.

Processing *C. elegans* for MIMS-EM imaging is similar to prior protocols for mammalian tissues^6^ with certain important adaptations. First, instead of conventional aldehyde-based fixation for EM applications, we vitrified live animals via high-pressure freezing (HPF) to preserve ultrastructure and coupled HPF with freeze-substitution^35,36^. This workflow overcomes the chemical fixation challenges created by the cuticle of *C. elegans*, which does not allow rapid infiltration of fixatives, while better preserving native membrane morphologies^35^. Following a more recent protocol originally developed for volume EM applications in *C. elegans*^36^, we incorporated ROTO (reduced osmium, thiocarbohydrazide and osmium) heavy-metal staining into the freeze substitution step to generate stronger membrane contrast for SEM. Finally, samples were embedded in resin blocks and sectioned onto silicon wafers for imaging.

In MIMS-EM, the target sample is first imaged using scanning electron microscopy (SEM) to map the organization and morphology of cells and their intracellular domains both at high resolution (5 nm) and across relatively large fields of view (**Figure 1C**). Next, the same sample is transferred to a MIMS microscope where a focused Cs^+^ primary ion beam is rastered across the section surface, generating secondary ions from the sample. The liberated secondary ions are separated by magnetic-sector mass spectrometry and detected using multiparallel detectors, ultimately providing simultaneous acquisition of ^12^C, ^13^C, ^14^N, ^15^N, ^32^S and ^31^P isotopes from spatially defined regions (i.e., pixels) of the sample^32,37^ (**Figure 1D**). Finally, the resulting images from both SEM and MIMS platforms are registered using fiducials and a custom algorithm to correct for imaging aberrations and artifacts, as previously established^6^. Using the registered MIMS-EM images, we can determine the relative rate of isotope turnover by creating isotope ratio maps (i.e., ^15^N-to-^14^N (^15/14^N) or ^13^C-to-^12^C (^13/12^C) ratios), which allows MIMS-EM to resolve metabolic turnover rates across spatial and temporal scales. Moreover, registration with SEM-derived ultrastructure enables us to assign the specific isotope tracers to the identities of the cells and organelles with which they are associated^6,7,37^ (**Figure 1E**).

We first used MIMS-EM to confirm the labeling efficiency of our stable isotope worm feeding approach. When adult animals were measured immediately following feeding (**Figure 1A**), we found ^15/14^N and ^13/12^C enrichment reaching levels >15,000x and >4,000x, respectively, in certain structures. Given that the natural ^15/14^N and ^13/12^C abundances fall around 0.00367 and 0.0112^37^, these ratios confirm efficient metabolic incorporation with ^13^C and ^15^N isotopes to near-saturating levels. Notably, these ratios exceed the enrichments achieved in prior mammalian studies by multiple orders of magnitude^6,7,38^, highlighting that experimental designs involving a large dynamic range of isotope levels may be a particular strength of the worm model. Next, we sought to determine the spatial resolution of our MIMS-EM approach, given that SEM and MIMS have a working maximum resolution of 5 nm and 50 nm, respectively. Here, we took advantage of a feature of cuticle ultrastructure clearly detectable in both SEM and MIMS images to estimate the effective resolution of our MIMS experiments (**Figure 1F**). By interpolating the full width at half-maximum (FWHM) of each peak, we found that 3.2 MIMS pixels equated to 36.7 SEM pixels. With SEM resolution set to 5 nm/pixel, our effective MIMS resolution is therefore estimated at ∼60 nm in X-Y (**Figure 1G**).

Upon initial examination, MIMS-EM ^13/12^C and ^15/14^N ratios reveal several expected tissue signatures. The cuticle, composed primarily of collagen, is re-synthesized at multiple larval molting stages during development, but is not thought to actively turnover between molts. We thus predicted these structures would develop particularly high ratios of heavy isotopes. Consistent with this prediction, MIMS-EM showed the highest heavy nitrogen incorporation in the cuticle with ^15/14^N ratios ∼15,000x (**Figure 1F**). Similarly, muscle myofilaments, composed of densely packed actin and myosin molecules, revealed high incorporation of labeled nitrogen with ^15/14^N ratios consistently >10,000 (**Figure 2A-C**). Lastly, we took advantage of the developmental differences between neuronal and hypodermal nuclei. Neuronal nuclei are born during embryonic development within the egg, prior to labeling with stable isotopes in our approach. In contrast, hypodermal nuclei undergo endoreduplication during larval development, thus incorporating dietary heavy isotopes. Confirming the expected labeling differences between embryonic and larval structures, we found that the neuronal nucleus had less than half the ^15/14^N ratios and ^13/12^C ratios of a hypodermal nucleus within the same animal cross-section (**Figure 2D-E**). Interestingly, most of the heavy isotope signal is seen with electron-dense areas within the nuclei, likely coinciding with heterochromatin (**Figure 2D**). The highest ratio for both ^15/14^N and ^13/12^C was in the nucleolus (**Figure 2E**), consistent with its role in ribosome biogenesis requiring ongoing incorporation of dietary resources throughout development. Collectively, these results validate predicted nutrient patterning while also highlighting the utility of MIMS-EM for resolving and quantifying turnover within the nuclear architecture^39,40^.

**Figure 2.**
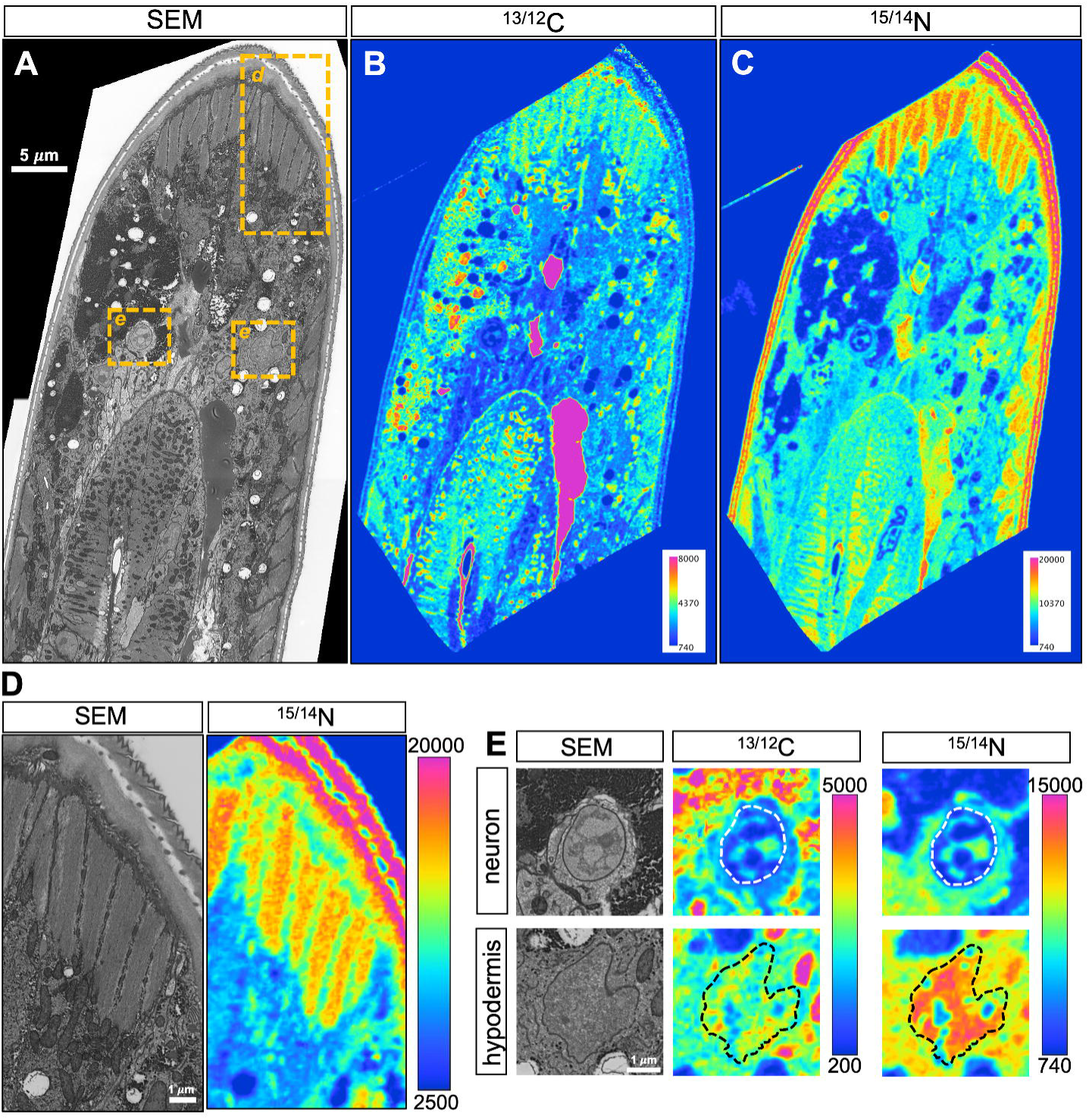
MIMS-EM resolves expected tissue and organelle isotope signatures. (A) SEM cross-section through the anterior region of a D1 animal labeled with ¹³C/¹ □ N from hatching. Boxed regions are shown in (D) and (E). Scale bar, 5 µm. (B) ^13/12^C ratio image of the section shown in (A). (C) ^15/14^N ratio image of the section shown in (A). (D) SEM and ^15/14^N ratio images of the cuticle and body wall muscle, showing high isotope in-corporation in the cuticle and myofilaments. Scale bar, 1 µm. (E) SEM, ^13/12^C, and ^15/14^N ratio images of a neuronal nucleus (top) and a hypodermal nucleus (bottom) from the same animal. Dashed outlines indicate nuclear boundaries. Scale bar, 1 µm.

### MIMS-EM reveals altered turnover at the tissue-scale during early-phase dietary restriction

Having established robust metabolic labeling and high-resolution structural imaging in *C. elegans*, we next deployed MIMS-EM to assess metabolic turnover across tissues and organelles of animals between distinct nutrient states. Dietary restriction (DR) robustly extends healthspan and lifespan across diverse species^8–10^, and evolutionary frameworks propose that these benefits arise from a reallocation of nutrient resources away from growth and reproduction and toward cellular maintenance and stress resilience^13^. At a coarse level, biochemical analyses comparing nutrient flux into somatic tissues vs. eggs in *Drosophila* have provided some support for this model, revealing that well-fed animals investment substantially more resources into progeny than animals undergoing DR^41^. However, the study also revealed that nutrient investments into the somatic tissues were higher in absolute terms in well-fed animals, indicating a need for more complex models of nutrient allocations than a simple two-compartment system (soma vs. egg) can provide. How such metabolic reorganization is coordinated across tissue and organelle scales within a single organism remains poorly resolved, but *C. elegans* MIMS-EM is uniquely suited to this question. In the worm it is possible to capture cross-sections representing virtually all tissue types of the organism at sufficient resolution to quantify isotope dynamics simultaneously across tissues and their contained organelles.

Because *C. elegans* allocate a large fraction of their nutrient resources to reproduction, typically producing hundreds of embryos in the first days of adulthood, the germline can dominate measurements of nutrient flux. To focus on how DR reallocates nutrients among somatic tissues, we therefore performed our analyses in a germline-deficient *glp-1* mutant background, effectively eliminating this reproductive sink. Because germline ablation can also independently extend lifespan^42,43^, we first tested and confirmed that the lifespan-extending effects of DR remain additive in *glp-1* mutants (**Figure S2**).

Eggs from *glp-1* mutants were placed on isotope-labeled bacterial lawns and raised to the first day of adulthood (D1). A subset of animals was harvested at D1 as a near-saturated baseline (t = 0), and the remainder were transferred to unlabeled bacterial lawns either concentrated for ad libitum (AL) feeding or diluted for dietary restriction (DR) using established methods^33^ (**Figure 3A**). Animals were collected at 8, 24, 48, and 72 h after the chase began, processed by high-pressure freezing and freeze-substitution, and sectioned through the anterior region of the animal. Sequential SEM and MIMS imaging of each section yielded co-registered structural and isotope ratio images across the tissues of *C. elegans* (**Figure 3B-D**). Manual segmentation of muscle, hypodermis, and intestine in each section enabled us to extract mean ^13/12^C and ^15/14^N ratios per tissue at each timepoint. We fit one-phase exponential decay models to these data and compared rate constants between dietary conditions using global nonlinear regression (**Figure 3E-I, Table S1**). At baseline under AL feeding, all three somatic tissues exhibited clear carbon turnover, with intestine displaying a shorter carbon half-life (t½ = 6.65 h) relative to muscle (12.7 h) and hypodermis (12.3 h) (**Figure 3H**). The elevated intestinal carbon turnover likely reflects the central role of this tissue in nutrient absorption and mobilization in *C. elegans*, integrating many functions of the vertebrate liver. Nitrogen half-lives (**Figure 4, Table S1**) were more similar across tissues (11.1–13.8 h) (**Figure 4H**) and consistently slightly longer than carbon, suggesting potential differences in the turnover rates of relatively carbon- vs. nitrogen-enriched macromolecular pools (e.g., membrane lipids vs. proteins).

**Figure 3.**
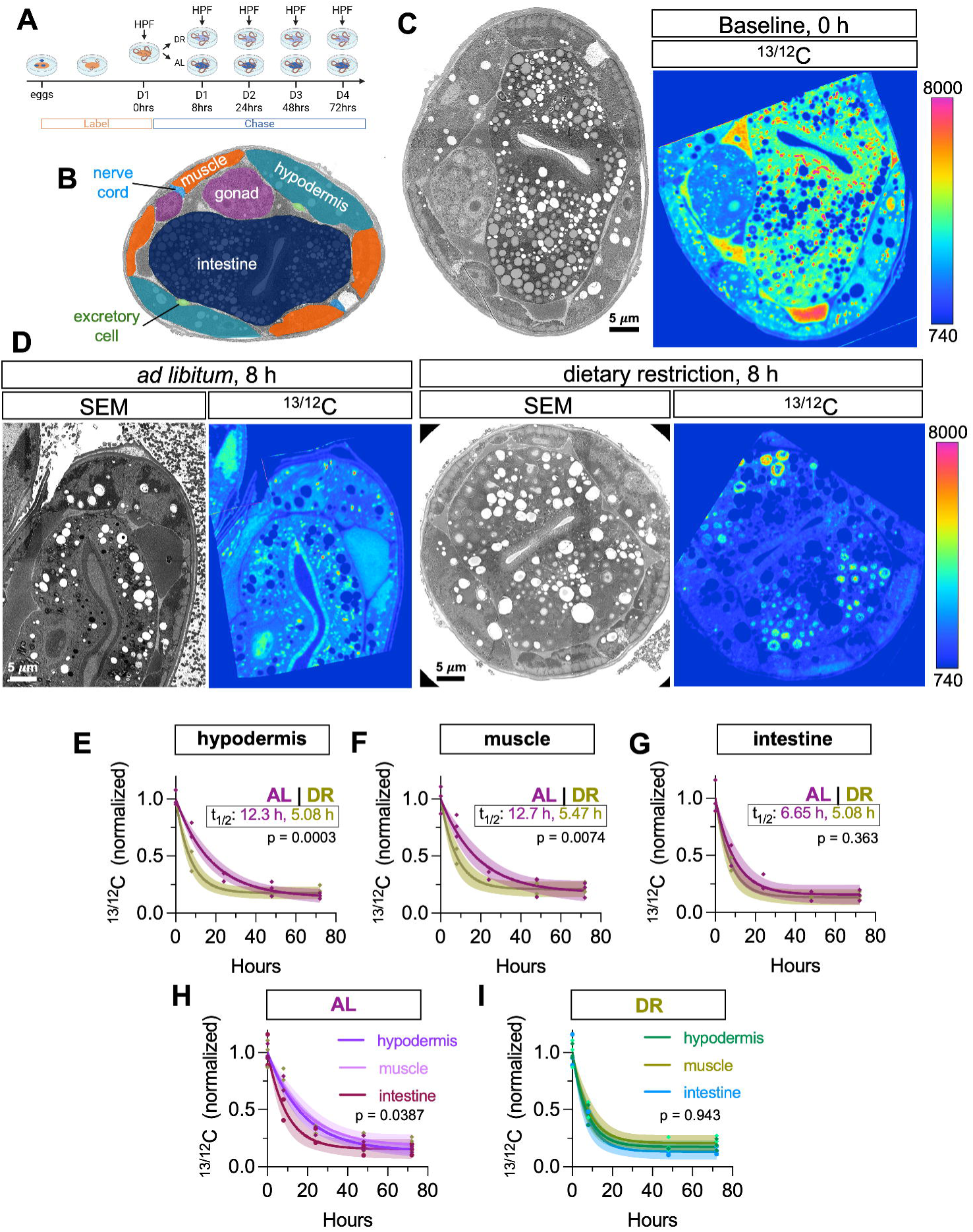
Dietary restriction accelerates ¹³/¹²C turnover in muscle and hypodermis but not intestine. (A) Pulse-chase design where *glp-1* animals were labeled with ^13^C/^15^N from hatching to D1, then transferred to unlabeled bacteria under AL or DR conditions and collected at the indicated chase durations. (B) Schematic of a *C. elegans* cross-section indicating the tissues analyzed. (C) SEM and ^13/12^C ratio images of a representative animal at baseline (0 h). Scale bar, 5 µm. (D) SEM and ^13/12^C ratio images of representative AL and DR animals after 8 h of chase. Scale bars, 5 µm. (E–G) ^13/12^C turnover in hypodermis (E), muscle (F), and intestine (G), normalized to the 0 h mean for each tissue. Each point is one animal (n = 2–3 animals per timepoint). Curves show one-phase exponential decay fits with 95% confidence bands. Half-lives are indicated; AL and DR decay rate constants were compared by extra sum-of-squares F test. (H, I) ^13/12^C turnover compared across tissues within AL (H) and DR (I) animals, plotted from the same data as (E–G). Decay rate constants were compared across tissues by extra sum-of-squares F test.

**Figure 4.**
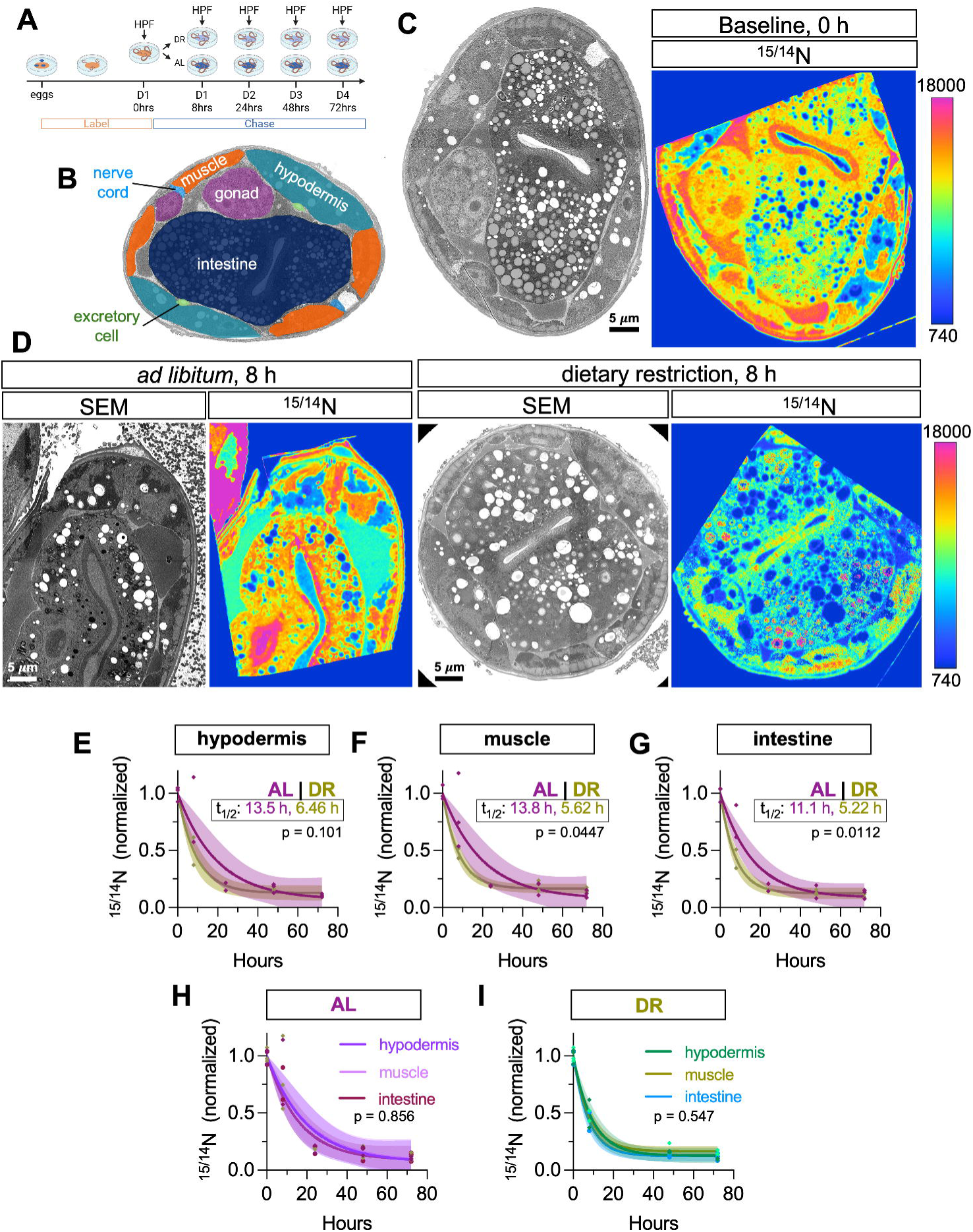
Dietary restriction accelerates ^15/14^N turnover across multiple tissues. (A) Pulse-chase design, as in Figure 3A. (B) Schematic of a *C. elegans* cross-section indicating the tissues analyzed. (C) SEM and ^15/14^N ratio images of a representative animal at baseline (0 h). Scale bar, 5 µm. (D) SEM and ^15/14^N ratio images of representative AL and DR animals after 8 h of chase. Scale bars, 5 µm. (E–G) ^15/14^N turnover in hypodermis (E), muscle (F), and intestine (G), normalized to the 0 h mean for each tissue. Each point is one animal (n = 2–3 animals per timepoint). Curves show one-phase exponential decay fits with 95% confidence bands. Half-lives are indicated; AL and DR decay rate constants were compared by extra sum-of-squares F test. (H, I) ^15/14^N turnover compared across tissues within AL (H) and DR (I) animals, plotted from the same data as (E–G). Decay rate constants were compared across tissues by extra sum-of-squares F test.

DR significantly accelerated carbon turnover in muscle (t½ 5.47 vs. 12.7 h; p = 0.0074) and hypodermis (t½ 5.08 vs. 12.3 h; p = 0.0003) but did not significantly alter intestinal carbon turnover (t½ 5.08 vs. 6.65 h; p = 0.363) (**Figure 3E-I**). The unchanged intestinal kinetics under DR are notable given the substantial acceleration observed in muscle and hypodermis and stem largely from the elevated baseline turnover of carbon in the intestine. On the other hand, nitrogen turnover responded to DR in a partially divergent pattern (**Figure 4**). In muscle, DR significantly accelerated nitrogen turnover (t½ 13.8 vs. 5.62 h; p = 0.0447), and a similar trend occurred in the hypodermis, but did not reach significance thresholds (t½ 13.5 to 6.46 h, p = 0.101). In contrast, DR significantly accelerated nitrogen turnover in intestine (t½ 5.22 vs. 11.1 h; p = 0.0112) (**Figure 4A-I**). Thus, the intestine is the only tissue in which DR significantly alters nitrogen but not carbon turnover. This dissociation between carbon and nitrogen kinetics suggests that DR does not uniformly accelerate macromolecular flux across tissues; rather, it tunes the turnover of distinct biomass classes in a tissue-specific manner. Such element-specific signatures represent a class of biological information that dual-element MIMS-EM is uniquely suited to capture.

Finally, we asked whether the turnover dynamics observed across the tissue extend to the mitochondrial networks within these respective cells. Segmenting individual mitochondria and extracting the mean carbon and nitrogen isotope ratios allowed us to fit the same one-phase decay models to the complete mitochondrial network within each tissue type (**Figure 5A**). Under AL conditions, the mitochondrial carbon turnover different significantly between tissue types (**Table S1**; p = 0.0058; extra sum-of-squares F test). Additionally, mitochondrial turnover was consistently faster than the corresponding whole-tissue turnover rates across all three tissues (hypodermis t½ = 7.23 vs. 12.3 h; muscle t½ = 10.6 vs. 12.7 h; intestine t½ = 4.87 vs. 6.65 h), while nitrogen half-lives were more comparable (hypodermis 12.9 vs. 13.5 h; muscle 13.6 vs. 13.8 h; intestine 12.3 vs. 11.1 h) (**Figures 3, 4, 5B-G, Table S1**). Similar to observations across the entire tissue areas, DR accelerated mitochondrial carbon turnover in hypodermis and muscle, but not the intestine (**Figure 5B-D**). Mitochondrial nitrogen turnover was significantly accelerated by DR in hypodermis (t½ 5.38 vs. 12.9 h; p = 0.0322) and intestine (t½ 5.38 vs. 12.3 h; p = 0.0348), with a non-significant trend in muscle (t½ 6.33 vs. 13.6 h; p = 0.0961). These results indicate that mitochondrial networks broadly track the tissue-specific and element-specific turnover patterns established above, while carbon in particular cycles through the mitochondria faster than through the tissue as a whole.

**Figure 5.**
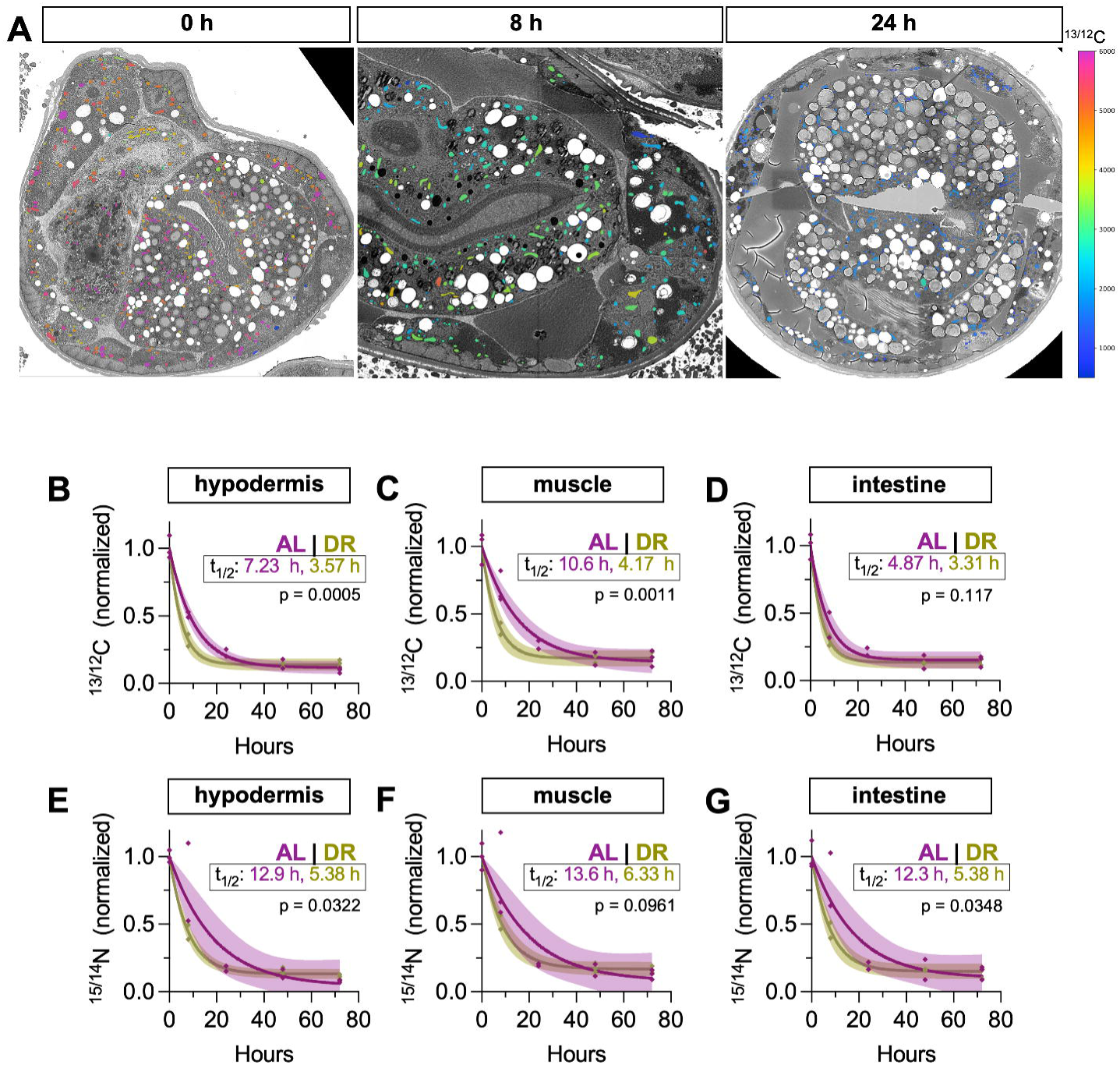
Mitochondrial turnover parallels tissue-level kinetics. (A) SEM images with segmented mitochondria each colored by its respective ^13/12^C ratio, shown for representative AL animals at 0, 8, and 24 h. Color scale is common to all three panels. (B-D) Mitochondrial ^13/12^C turnover in hypodermis (B), muscle (C), and intestine (D), normalized to the 0 h mean for each tissue. Each point is one animal (n = 2–3 animals per timepoint). Curves show one-phase exponential decay fits with 95% confidence bands. Half-lives are indicated; AL and DR decay rate constants were compared by extra sum-of-squares F test. (E-G) Mitochondrial ^15/14^N turnover in hypodermis (E), muscle (F), and intestine (G), plotted and analyzed as in (B-D).

### Mitochondrial heterogeneity is constrained and spatially organized

The single-organelle resolution of MIMS-EM enabled a direct test of how heterogeneity is structured within mitochondrial populations *in situ*. Mitochondrial networks are continuously remodeled by fission and fusion, processes that can equilibrate organelle content and buffer composition across the network^16,44^. In certain cell types, however, such as neurons and skeletal muscle cells, some individual mitochondria appear to be segregated from the network and remarkably long-lived^38,45^. The extent to which a mitochondrial network within a cell behaves as a coordinated, equilibrating population or as a collection of independent subtypes with distinct turnover rates remains debated and likely context-dependent^46^.

To address this, we examined the distribution and spatial organization of ^13/12^C ratios of thousands of individual mitochondria across the tissues and conditions of our dataset. First, we established a baseline measure of mitochondrial-scale heterogeneity by computing the coefficient of variation (CoV) of ^13/12^C ratios across all mitochondria within each animal. Because CoV normalizes variance to the population mean, this metric accounts for the ∼10-fold decline in absolute labeling that occurs over the chase period. We found that mitochondrial CoV remained remarkably stable across the time course (**Figure 6A**), with mean values between 0.22 and 0.27 maintained over 72 h. In AL animals, CoV varied only modestly by tissue (p < 0.045, two-way ANOVA) with no pairwise comparisons reaching statistical significance (**Figure 6B**). Lastly, although DR generally appeared to reduce inter-individual variation in tissue turnover (**Figure 3I, 4I**), diet had no discernable effect on mitochondrial heterogeneity (**Figure 6C**). To exclude the possibility that isotope variation at the mitochondrial level reflects measurement noise rather than biological heterogeneity, we performed a split-half analysis of ^13/12^C signal within individual mitochondria. The within-organelle measurement error (median ∼0.002 of the population mean) was ∼95-fold smaller than the dispersion between organelles, supporting that the observed CoV reflects biological heterogeneity between mitochondria (**Table S2**).

**Figure 6.**
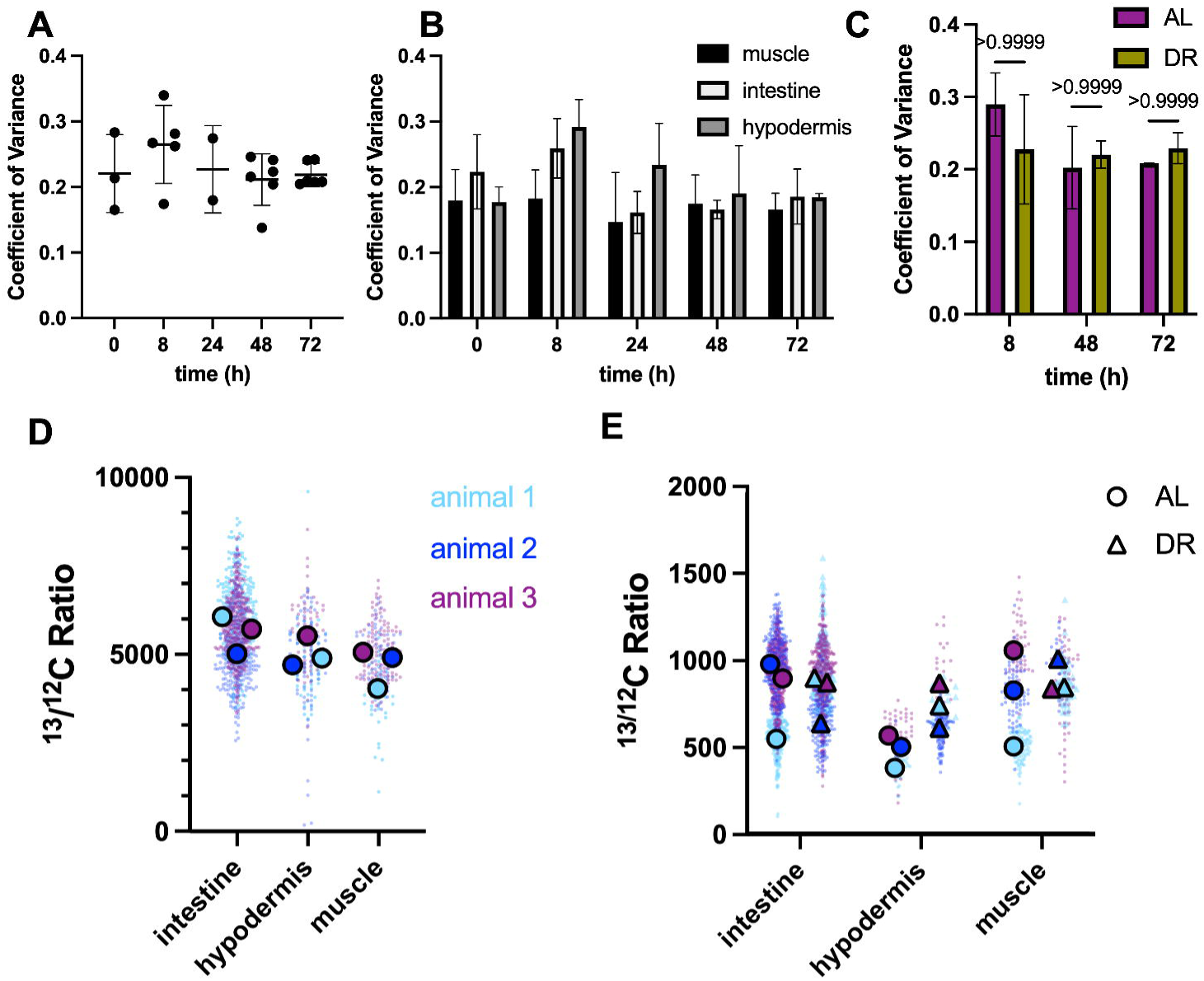
Mitochondrial ^13/12^C heterogeneity is maintained within a narrow range. (A) CoV of mitochondrial ^13/12^C ratios within each animal, pooled across all tissues and combining ad libitum (AL) and dietary-restricted (DR) animals, plotted against chase duration. Each point is one animal (n = 3, 5, 2, 6, and 6 animals at 0, 8, 24, 48, and 72 h, respectively). Plotted as mean ± SD, analyzed as one-way ANOVA: p = 0.38 and no pairwise comparison reached significance (Tukey). (B) CoV of mitochondrial ^13/12^C ratios in AL animals, separated by tissue. Bars show mean ± SD across animals; n = 3 everywhere except muscle (24 h) and intestine (48 h), n = 2. Analyzed by two-way ANOVA: p = 0.026 (time), p = 0.045 (tissue), and no pairwise comparison reached significance (Sidak). (C) CoV of mitochondrial ^13/12^C ratios by diet at matched chase timepoints. Bars show mean ± SD across animals; n = 3 for all conditions except 8 h DR (n = 2). Analyzed by two-way ANOVA: p = 0.13 (time), p = 0.86 (diet), and pairwise comparisons shown on graph (Tukey). (D) Distribution of ^13/12^C ratios of individual mitochondria at 0 h by tissue and animal. Small dots represent individual mitochondria; large outlined dots represent per-animal means. (E) Distribution of ^13/12^C ratios of individual mitochondria at 72 h, by tissue and diet. Small dots represent individual mitochondria, colored by animal as in (D); large outlined dots represent per-animal means (AL, circles; DR, triangles).

If discrete subpopulations of mitochondria were uncoupled from the general turnover dynamics of the cell, we would expect to observe outliers or multimodal distributions emerge during the chase. To assess this, we examined per-mitochondrion ^13/12^C distributions at t = 0 and t = 72 h, where mitochondria refractory to turnover would be most apparent (**Figure 6D-E**). At baseline, distributions were tight and centered near saturation across all three tissues. At 72 h, the distributions remained unimodal and relatively continuous, with little evidence for discrete sub-populations that retained label. These data support a model in which the network as a whole undergoes equilibration in these cell types and conditions, or that turnover processes are applied relatively uniformly across all individual mitochondria in the network. Future studies involving direct modulation of mitochondrial fission/fusion could limit inter-mitochondrial mixing and distinguish between these possibilities.

To test more directly whether mitochondrial labeling is spatially organized within tissues, we incorporated positional information into our analysis. As a simple proof-of-principle, we can identify two adjacent muscle cells in a cross-section and compare the isotope ratios of their mitochondrial networks on the basis of position (**Figure 7A-B**). These distinct mitochondrial populations exhibited clear ^13/12^C differences (p < 0.0001), confirming that MIMS-EM can resolve metabolic distinctions between distinct cells that would otherwise remain obscured in bulk biochemical measurements.

**Figure 7.**
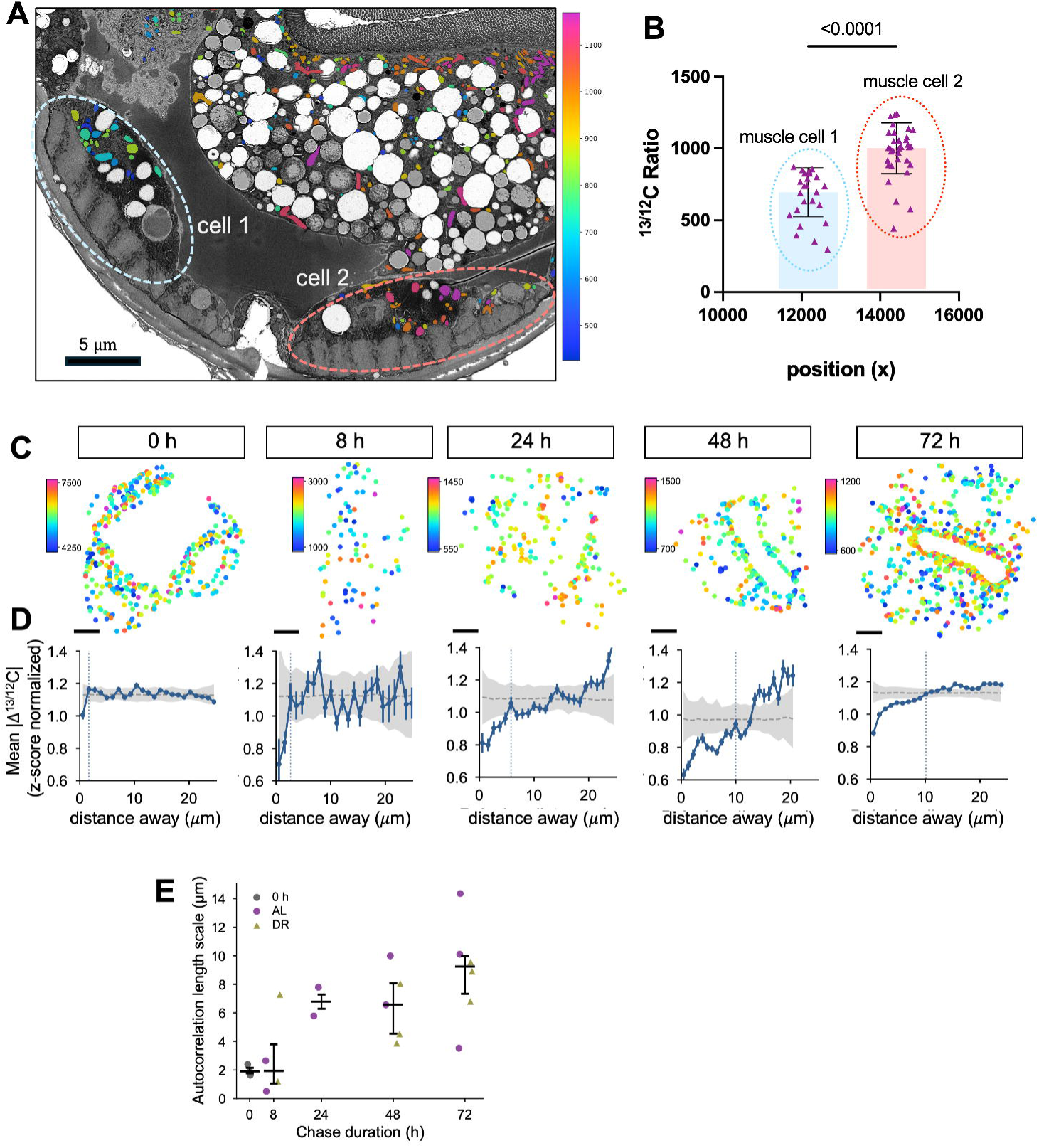
Mitochondrial ^13/12^C labeling is spatially autocorrelated and expands over time. (A) SEM cross-section showing two adjacent muscle cells (dashed outlines) from a DR animal at 72 h (827_3), with segmented mitochondria colored by ^13/12^C ratio. Scale bar, 5 µm. (B) ^13/12^C ratios of individual mitochondria from the two muscle cells shown in (A), plotted against position along the section x axis. Each point is one mitochondrion (n = 25 and 34 for cells 1 and 2); bars show mean ± SD. Analyzed by Welch’s t test. (C) Centroid positions of intestinal mitochondria from a representative AL animal at each chase timepoint colored by ^13/12^C ratio. Color scales are set independently for each animal. Scale bars, 5 µm. (D) Variograms for the animals shown in (C), plotting the mean absolute difference in z-scored ^13/12^C ratio between pairs of mitochondria against the distance separating them. Points show mean ± SEM per distance bin; gray band and dashed line show the 95% range and mean of the permuted null distribution; dotted vertical line marks the autocorrelation length scale. Per-animal Moran’s I, permutation P values, and length scales are reported in Table S3. (E) Autocorrelation length scale by chase duration. Each point is one animal (n = 20), with AL and DR combined for analysis; bars show median ± interquartile range. Analyzed by Spearman rank correlation: ρ = 0.72, p = 0.0003.

We next applied this positional framework to the larger, intracellular architecture of the intestinal cells. First, we mapped the centroid positions and isotope ratios of individual mitochondria over the AL time course (**Figure 7C**). We then analyzed all pairwise combinations of mitochondria within each of these animals, assessing their distance and the similarity between their ^13^C isotope signatures, normalizing isotope ratios by z-score to account for differences in absolute labeling levels across timepoints (**Figure 7D**). This assessment of spatial autocorrelation within each of these animals confirmed that neighboring mitochondria are more similar to one another in terms of isotope composition than distant mitochondria (**Figure 7D**; **Table S3;** statistically significant in 19 of 20 animals by Moran’s I). This local similarity was apparent across animals under both AL and DR conditions (**Table S3**). Notably, the distance at which neighboring mitochondria correlated expanded over the timecourse of the chase, from <2 µm at t = 0 to ∼9 µm by 72 h (**Figure 7D, E**; Spearman ρ = 0.72, p = 0.0003). This result indicates that local similarity emerges and expands as turnover proceeds, consistent with a model of localized mixing of mitochondrial contents that progressively equilibrates across the cell’s larger mitochondrial network. Collectively these results highlight the ability of MIMS-EM to detect organelle-level heterogeneity and subsequently apply spatial context and morphometry to develop mechanistic hypotheses underlying how cells organize and regulate organellar biogenesis and turnover.

## Discussion

The compact size and simple body plan of *C. elegans* offer the potential to capture metabolic turnover across the tissues of an intact metazoan at sub-organelle resolution. To realize this potential, here we adapted MIMS-EM for the worm. We developed metabolic labeling strategies, sample preparation for correlative MIMS and SEM imaging, and analytical approaches for quantifying isotope turnover, then applied this workflow to assess how animals adapt to the early stages of dietary restriction. Across cellular and organelle scales, we identified diet- and element-specific effects. Different tissues displayed distinct turnover kinetics under both AL and DR diets, turnover of carbon and nitrogen pools was partly uncoupled, and heterogeneity within a given cell’s mitochondrial network was revealed. Through the application of MIMS-EM to *C. elegans,* these observations begin to reveal the principles by which an intact animal allocates nutrients across its complement of distinct tissues, cells, and organelles.

Our findings also impact current mechanistic models for how DR extends lifespan. Based on its tendency to activate autophagic processes, DR is often presumed to promote health in part by increasing the turnover and bulk recycling of cellular material^27,28^. Yet studies that directly measure proteome turnover in liver and muscle cells have repeatedly yielded contradictory results, showing instead that a greater number of proteins exhibit slowed turnover than accelerated turnover during DR^29–31^. These findings suggest instead that DR rebalances biosynthesis and degradation of certain protein classes through a more selective program. Our MIMS-EM measurements show that DR enhances overall turnover of carbon and nitrogen biomass, which might seem to support the general recycling-based model. Importantly, however, these approaches quantify turnover differently. Tandem mass spectrometry resolves the kinetics of individual peptides and proteins, but achieves only partial coverage of the proteome and reports each protein’s turnover rate independently of how much that protein contributes to the total. An elemental measurement instead reflects the summed flux of carbon and nitrogen across the entire macromolecular pool. Because a smaller number of highly abundant proteins can contribute disproportionately to this total elemental pool, elemental turnover can therefore rise even as the turnover rate of many individual proteins slows. Our data thus point to the possibility that these competing models are not mutually exclusive; global turnover can be enhanced at the level of elemental accounting, while cells can still accomplish protein- or organelle-specific functional remodeling.

From our analysis of turnover at the tissue scale, the intestine illustrates why models for DR effects on turnover should not be assumed to extrapolate across tissue types. Specifically, the intestine exhibited higher baseline turnover of carbon compared to muscle and hypodermis, and no effect of DR, contrasting with the substantial acceleration observed in the other tissues. The accelerated baseline turnover patterns likely reflect the intestine’s role as the hub of nutrient uptake and distribution in the worm. However, nitrogen pools behaved differently. Intestinal nitrogen turnover was comparable between tissues at baseline and significantly accelerated by DR. This apparent uncoupling between carbon and nitrogen kinetics at the tissue level suggests differential handling of lipid (carbon-rich and nitrogen-poor) and protein (both carbon- and nitrogen-rich) macromolecular pools. While MIMS-EM is well-suited to generating hypotheses regarding these distinct macromolecular classes, resolving them requires additional approaches. For example, coupling MIMS-EM with parallel, biochemical analysis of purified macromolecular pools would allow carbon and nitrogen signals to be assigned to specific molecules and molecular classes^45^, providing the molecular identities that correspond with the spatial measurements. Additionally, future MIMS-EM experiments may utilize isotope-labeled precursors specific to distinct macromolecular classes (e.g., labeled amino acids for incorporation into proteins).

The single-organelle resolution provided by MIMS-EM allowed us to examine turnover rates across the mitochondrial network contained within individual cells. Across tissues, timepoints, and diets, the variability in inter-mitochondrial isotope content remained largely fixed. Over the three days of the timecourse, which notably allowed isotope decay curves to reach a plateau phase, we observed little evidence of outlier subpopulations of mitochondria that were refractory to turnover. Combined with our finding that neighboring mitochondria were generally more similar than distant mitochondria in a cell, these observations are consistent with the dynamic connectivity of mitochondrial networks, continually mixing contents through fission and fusion rather than existing as separate, uncoupled organelles. Performing MIMS-EM analysis while manipulating fission/fusion processes will provide more direct test of this model. While our analysis of mitochondrial heterogeneity focused on the intestine, it is important to note that the connectivity of mitochondrial networks increasingly appears to be cell-type specific. Key examples include cardiac muscle and certain neurons, which respectively exhibit less coupling generally or in specific subcellular regions^7,46^. Given the increased heterogeneity that emerges with aging across molecular, organellar and cellular scales^7,47–49^, future MIMS-EM studies may be useful in mapping when and where this heterogeneity arises. Moving forward, *C. elegans* provides a powerful pairing with mammalian models for elucidating the mechanisms underlying age-onset metabolic heterogeneity as well as the consequences for tissue function and organismal healthspan.

When coupled with pulse-chase labeling, MIMS-EM brings metabolic reallocation from a conceptual framework to directly measurable observations. The capacity to visualize multiple isotope labels in parallel further allows the fates of distinct nutrient types to be followed in parallel within the same specimen. Combined with the compact body plan and experimental tractability of *C. elegans*, MIMS-EM offers a way to follow how genetic, dietary and age-related changes reshape metabolic fates from the organelle to the organismal scale, generating remarkably rich datasets that can support integrative models of animal physiology in diverse, native contexts.

## Methods

### *C. elegans* strains and maintenance

*C. elegans* N2 Bristol and *glp-1(e2141)* III (strain CB4037) strains were obtained from the *Caenorhabditis* Genetics Center maintained at 20 °C on standard nematode growth media (NGM) plates seeded with E. coli OP50. To compare worm growth between labeled and unlabeled media, N2 adult worms were allowed to lay eggs for 2 h and eggs were picked onto labeled or unlabeled plates. On day 1 of adulthood, N2 worms were anesthetized with 1 mM of tetramisole (Fisher Scientific, 50-004-33485) and imaged. Worm size was calculated through ImageJ software (ImageJ 1.54f). For *in vivo* isotope labeling, N2 worms were bleached onto isotope-labeled MM plates, allowed to grow to day 1 adults at 20 °C and processed for electron microscopy. To assess isotope turnover, synchronized eggs were allowed to develop on isotope-labeled MM plates until day 1 adulthood. From the same population, a subset was processed for HPF at time = 0 and remaining worms were picked onto unlabeled AL or DR plates and sub-populations were again processed for HPF 8, 24, 48 and 72 h after plating. To impair germline formation, temperature-sensitive *glp-1* mutant worms were bleached onto labeled MM plates and incubated at 25 °C for 40 h before being shifted to standard 20 °C for 22 h, when worms reached day 1 adulthood.

### Lifespan Analysis

Timed egg lays were used to obtain synchronized populations. One day before adulthood (day 0), at least 100 mid-L4 worms were transferred to 5 or 10 fresh plates at a density of 10 or 20 worms per plate. To separate from progeny and avoid starvation, worms were transferred to fresh plates every day or every other day until the first deaths (∼10-12 days). Survival was scored every 1-2 days and worms were marked dead when they failed to respond to three taps on the head and tail. Worms were censored for leaving the plate and desiccating and for reproduction-related deaths (internal hatching and loss of vulval integrity). Logrank Mantel-Cox tests were performed to determine statistical significance.

### Isotope labeling and plate preparation

Minimal media (MM) is composed of M9 buffer supplemented with 22.2 mM of labeled [U-^13^C_6_] glucose (Cambridge Isotope Laboratories, Inc., CLM-1396), 18.7 mM of labeled ^15^NH_4_Cl (Cambridge Isotope Laboratories, Inc., NLM-467), and 0.178 mM uracil (Sigma-Aldrich, U0750). Minimal media was filtered through 0.22 µm pore size filter. Ad libitum (AL) and dietary restricted (DR) plates were prepared following Ching et al.^33^. Briefly, an individual *E. coli* OP-50 colony was picked and grown in labeled minimal media (MM) or Isogro growth media (CM) (Sigma-Aldrich, 606839) resuspended at 1 mg/mL in labeled MM media for 8 h at 37 °C. To grow a large volume, this culture was inoculated at 1:130 dilution and grown overnight for 16 h at 37 °C. The bacterial concentration was calculated by measuring OD600 through a NanoDrop 2000c. The culture was centrifugated at 12000 rcf for 15 min and resuspended at 2 ×10^10^, 10^8^ and 10^11^ bacterial concentrations for maintenance bacterial lawns, DR lawns, and AL lawns, respectively. Bacterial suspensions were treated with Carbenicillin and Kanamycin at final concentrations of 5 mg/mL and 2.5 mg/mL, respectively, before plating 200 µL per NGM plate.

To compare labeled and unlabeled media, unlabeled glucose (Sigma Aldrich, G7528) and ammonium chloride (Fisher Scientific, A661-500) were utilized. After 16 hr incubation, the OD600 of the bacterial culture was measured through a NanoDrop 2000c. Three bacterial concentrations were prepared; 6.5 × 10^9^ for bacterial concentration was plated and allowed to dry for 48 hrs at room temperature, after which were stored at 4 °C.

### Sample processing for MIMS-EM

For high-pressure freezing, 0.15 M sucrose in M9 buffer was pipetted in the 200 uL deep side of an A-type carrier (Ted Pella, 39200). Worms were anesthetized in 1 mM of tetramisole (Fisher Scientific, 50-004-33485) for 5min and picked onto the A carrier. The assembly was covered by the flat side of a B-type carrier (Ted Pella, 39201) and vitrified using a high-pressure freezer (Leica EM ICE). Samples remained submerged in liquid nitrogen until further processing. Freeze substitution was carried out following Belanger et al.^36^. Briefly, frozen carriers were quickly placed in dry ice-chilled cryovials with 1% osmium tetroxide in acetone. Cryovials were kept in a Styrofoam box with dry ice at −20 °C overnight. Dry ice was then removed and cryovials were kept at −20 °C until the next day. Samples were then kept at 4 °C. Samples then underwent multiple incubations with decreasing concentration of acetone in double-distilled water and equilibrated in 0.1M sodium cacodylate buffer. Post-fixation was performed using a reduced osmium protocol with 1% osmium tetroxide, 0.75% potassium ferrocyanide in 0.1M sodium cacodylate. Samples were then stained with heavy metals with 1% thiocarbohydrazide, 2% osmium tetroxide and 1% uranyl acetate. Lastly, samples were dehydrated through a series of graded acetone until 100% acetone, prior to Epon812 resin infiltration. Infiltration was carried out by incubating samples in progressively increasing concentrations of Epon812 resin in acetone until multiple incubations in 100% Epon812 resin. After resin infiltration, worms were manually embedded in resin blocks and hardened at 60 °C for 72 hrs. Blocks were then thick sectioned with a glass blade until a desired location and 80 nm sections were collected with a diamond blade on SEM wafers (Electron Microscopy Services, cat# 71893-10). Wafers were then post-stained with 2% uranyl acetate and lead citrate.

### Correlative MIMS-EM image acquisition

Sections were mapped using SEM (Crossbeam 550, Zeiss, Germany). User-supervised image acquisition was guided using automated tile acquisition and image mosaicking software (Atlas 5, Fibics, Ottawa, Canada). Images were acquired with a pixel size of 5 nm across a complete animal cross-section. Next, wafers containing the mapped samples were transferred to a MIMS microscope (50 L NanoSIMS, Cameca, France) for acquisition of multi-isotope maps (^13^C, ^12^C, ^32^S, ^14^N, and ^31^P) as previously established^5,6^ using the following MIMS image acquisition parameters: image size of 512×512 pixels, raster size of 30-to-40 um^2^, at least three frames per raster with a 10 min acquisition time per frame using the beam adaptor D1-3.

### Image Registration and Processing

Multi-isotope imaging mass spectrometry (MIMS) data were acquired as multi-plane secondary-ion count images in Cameca .im files, which were read with the open-source sims Python library. For each ion species, the sequential acquisition planes were first corrected for inter-plane drift by frame-to-frame affine registration using the pystackreg Python library with StackReg in AFFINE mode and then summed along the z axis to yield a single accumulated count image per isotope. Each MIMS field of view was registered to its corresponding EM image. A user selected a set of corresponding landmarks on the EM image and on a reference secondary-ion image (typically ³²S, ¹²C, or ¹²C¹ N, whichever gave the clearest references for morphological similarities). From these landmark pairs a similarity transform was estimated (scikit-image), and a horizontal mirror was tested and applied when it reduced the landmark residual, giving a coarse rotation/flip alignment. Residual translational bias was removed using the median landmark offset, and remaining local, non-linear distortion was corrected with a thin-plate-spline (scikit-image) transform fit to the landmark displacements. Each summed isotope image was then resampled into EM/canvas space by applying the similarity transform followed by the TPS warp, and its transformed bounding box was recorded. The registered per-field isotope tiles were stitched into canvas-sized mosaics by cropped placement at their registered bounding boxes without blending, with earlier acquired acquisitions superseding at pixels of overlap. Isotope-ratio mosaics were formed by pixel-wise division of the corresponding stitched numerator and denominator mosaics (scaled ×10,000, with zero denominators set to one), producing the final registered isotope and isotope-ratio images used for downstream quantification and visualization. Individual tissue and mitochondrial boundaries were segmented manually from SEM images alone. ¹³C/¹²C and ¹ N/¹ N ratios within each segmented compartment were extracted from registered MIMS images. The sample inventory from this analysis is compiled in Table S4.

To calibrate MIMS pixel size, we measured line profiles of a shared cuticle structure in SEM and MIMS images, then used linear interpolation to identify the pixel values corresponding to half-maximum for each edge of the structure. From these, we calculated FWHM in each channel and generated a SEM:MIMS pixel ratio to estimate effective MIMS resolution.

### Statistical Analysis

For bacterial yield comparisons, when comparing two samples with one variable, an unpaired two-tailed *t* test was used. For more than two samples with one variable, a one-way ANOVA followed by a Dunnett’s multiple comparison test was used. A *p* value less than 0.05 was considered statistically significant and denoted as follows: *<0.05, **<0.01, ***<0.001 and ****<0.0001.

For isotope turnover curves, isotope ratios across the timecourse were normalized to 0 h, then fit by nonlinear regression using a one-phase exponential decay model (GraphPad Prism v11.0.0). Initial values were constrained to 1.0, whereas plateau and decay rate constant *k* were fitted for each condition. Differences is turnover kinetics were assessed via the extra sum-of-squares F test focusing on decay rate constant. Reported half-lives were calculated from fitted decay constants (t_1/2_ = ln 2 / *k*). Data are displayed as best-fit curve with 95% confidence interval.

To analyze mitochondrial heterogeneity, analyses were performed per tissue unless otherwise noted and restricted to animals with at least five segmented mitochondria in the given tissue. Coefficient of variation was calculated from all mitochondria within each tissue and reported per animal. To confirm that measured variation between individual mitochondria reflected biological differences rather than noise, a split-half analysis was performed on a representative section. For each mitochondrion, pixels within the segmented mask were randomly divided into two halves, the mean of each ratio was calculated, and the absolute difference between halves was normalized to the mean of the whole mitochondrion. The median of this within-mitochondrion difference was compared to the coefficient of variation measured between mitochondria.

Spatial analyses were performed in the intestine, which forms a continuous cellular field and contains higher numbers of mitochondria needed for analysis. For each animal, centroid positions and mean ^13/12^C ratios were used to assess spatial correlations. Autocorrelation was quantified as Moran’s I^50^ using inverse-distance weights and standardized by row to sum to one. Expected values, standard deviation and significance were determined from 500 random permutations of ratios among centroid positions. The spatial scale of autocorrelation was measured by variogram with pairwise distances between mitochondria binned into 24 equal-width bins spanning to the 95^th^ percentile. Mean and standard error were computed per bin, with bins containing fewer than 10 pairs excluded. A null distribution was generated by 200 permutations of ratios among positions, and the autocorrelation length was defined as the shortest distance at which the variogram entered the 95% range of the null distribution. To compare spatial structures across individual animals and timepoints, ratios were standardized by z-score within each animal. The relationship between length scale and chase duration was tested by Spearman rank correlation across all animals, with AL and DR animals combined. Analyses were performed in GraphPad Prism (v11.0.0) and Python (v3.11) using NumPy, SciPy, and pandas.

## Data and Code Availability

All source data will be published with the manuscript and large image files will be placed in a public generalist repository for community data-mining and analysis upon acceptance. Custom code for registration and extraction of MIMS-EM features is available through GitHub (https://github.com/ArrojoDrigoLab/MIMS-EM), and custom scripts used for standard statistical analyses and data plots are available on request.

## Acknowledgements

We thank all members of the Burkewitz and Drigo labs for helpful conversations, S. Sviben for technical advice, and V. Gama and C. Wright for valuable feedback. Electron microscopy sample preparation and imaging were performed in part through the use of the Vanderbilt Cell Imaging Shared Resource (CISR) supported by NIH grants CA68485, DK20593, DK58404, DK59637, EY08126, R24OD037694, S10MH137068, and S10OD028704. Electron microscopy consultation and services were also performed in the Harvard Medical School (HMS) Electron Microscopy Facility. We thank M. Ericsson (HMS), E. Krystofiak, M. Vinogradova and R. Hart (Vanderbilt CISR) for electron microscopy services. MIMS imaging was performed by Dr. Yunbin Guan at the Division of Geological and Planetary Sciences at Caltech. This work was supported by NIH/NIA grant R56AG082758 (KB and RAeD) and NIH/NIDDK T32DK07563 (CA).

---

**Supplemental Figure 1.**
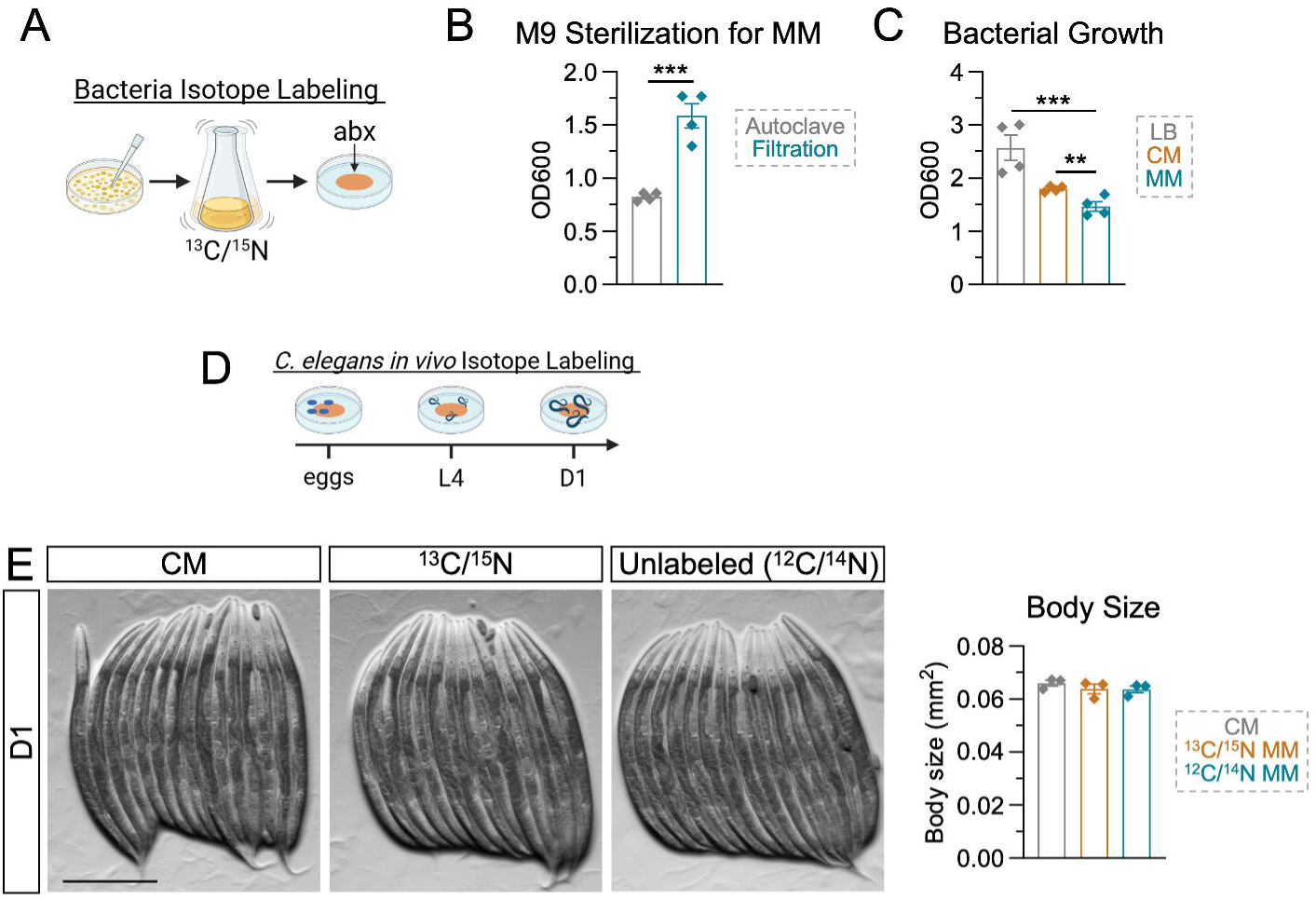
Isotope labeling of the bacterial diet does not affect *C. elegans* growth. (A) Scheme for generating ¹³C/¹ N-labeled bacterial lawns. (B) Bacterial yield in minimal media (MM) prepared with autoclaved versus filter-sterilized M9 buffer, measured as OD600. Each point is an independent culture (n = 4); bars show mean ± SEM. Analyzed by unpaired two-tailed t test: ***p < 0.001. (C) Bacterial yield in LB, complete media (CM), and minimal media (MM), measured as OD600. Each point is an independent culture (n = 4); bars show mean ± SEM. Analyzed by one-way ANOVA with Dunnett’s multiple comparisons test: **p < 0.01, ***p < 0.001. (D) Scheme for *in vivo* isotope labeling of *C. elegans*. (E) Representative images of D1 animals grown on bacteria cultured in CM, ¹³C/¹ N-labeled MM, or unlabeled MM, with quantification of body size. Each point is an independent biological replicate (n = 3 means each from a group of 10-15 animals); bars show mean ± SEM. Scale bar, 250 µm.

**Supplemental Figure 2.**
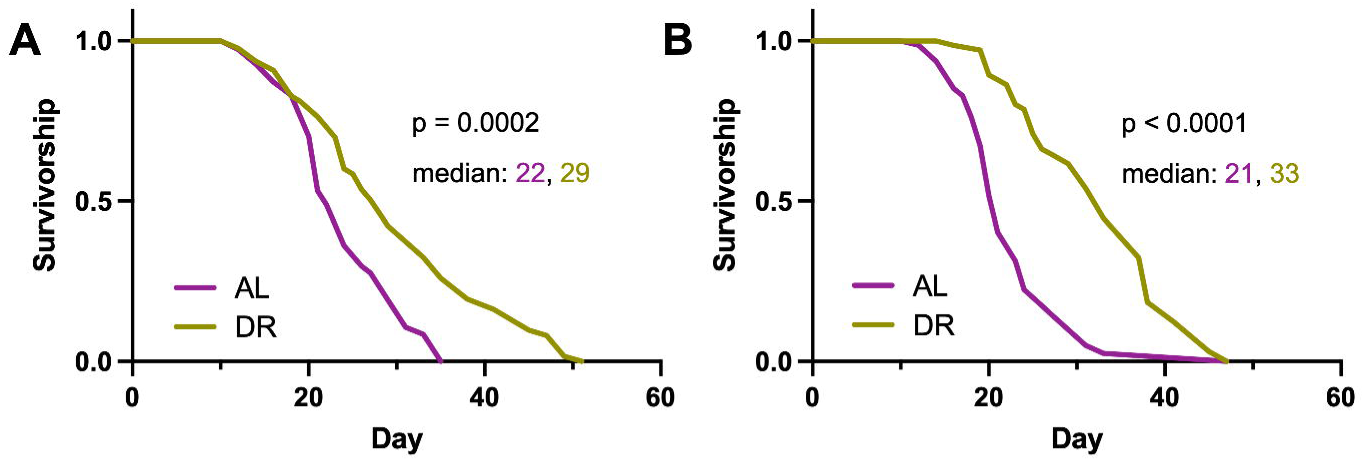
DR extends lifespan of *glp-1* mutants. (A, B) Independent survival analyses of *glp-1* mutants on either AL or DR plates made from MM-grown bacterial cultures. Starting n = 100 animals; p-values shown are derived from Logrank (Mantel-Cox) test.

**Table S1. Isotope decay curve statistics.**

**Table S2. Split-half Analysis.**

**Table S3. Spatial analysis of mitochondrial** ^13^**^/12^C signatures**

**Table S4. Inventory of MIMS-EM samples.**

## Bibliography

1. Finkel, T. (2015). The metabolic regulation of aging. Nat. Med. 21, 1416–1423.

2. Smith, H.J., Sharma, A., and Mair, W.B. (2020). Metabolic Communication and Healthy Aging: Where Should We Focus Our Energy? Dev. Cell 54, 196–211.

3. Jain, A., and Zoncu, R. (2022). Organelle transporters and inter-organelle communication as drivers of metabolic regulation and cellular homeostasis. Mol Metab 60, 101481.

4. Voeltz, G.K., Sawyer, E.M., Hajnóczky, G., and Prinz, W.A. (2024). Making the connection: How membrane contact sites have changed our view of organelle biology. Cell 187, 257–270.

5. Steinhauser, M.L., Bailey, A.P., Senyo, S.E., Guillermier, C., Perlstein, T.S., Gould, A.P., Lee, R.T., and Lechene, C.P. (2012). Multi-isotope imaging mass spectrometry quantifies stem cell division and metabolism. Nature 481, 516–519.

6. Habashy, A., Acree, C., Kim, K.-Y., Zahraei, A., Dufresne, M., Phan, S., Cutler, M., Patterson, E., Mulligan, A.G., Burkewitz, K., et al. (2025). Spatial patterns of hepatocyte glucose flux revealed by stable isotope tracing and multi-scale microscopy. Nat. Commun. 16, 5850.

7. Arrojo E Drigo, R., Lev-Ram, V., Tyagi, S., Ramachandra, R., Deerinck, T., Bushong, E., Phan, S., Orphan, V., Lechene, C., Ellisman, M.H., et al. (2019). Age Mosaicism across Multiple Scales in Adult Tissues. Cell Metab. 30, 343–351.e3.

8. Mair, W., and Dillin, A. (2008). Aging and survival: the genetics of life span extension by dietary restriction. Annu. Rev. Biochem. 77, 727–754.

9. Fontana, L., and Partridge, L. (2015). Promoting health and longevity through diet: from model organisms to humans. Cell 161, 106–118.

10. Green, C.L., Lamming, D.W., and Fontana, L. (2022). Molecular mechanisms of dietary restriction promoting health and longevity. Nat. Rev. Mol. Cell Biol. 23, 56–73.

11. Anderson, R.M., and Weindruch, R. (2010). Metabolic reprogramming, caloric restriction and aging. Trends Endocrinol. Metab. 21, 134–141.

12. Kirkwood, T.B., and Holliday, R. (1979). The evolution of ageing and longevity. Proc. R. Soc. Lond. B Biol. Sci. 205, 531–546.

13. Kirkwood, T.B.L. (2002). Evolution of ageing. Mech. Ageing Dev. 123, 737–745.

14. Gomes, L.C., Di Benedetto, G., and Scorrano, L. (2011). During autophagy mitochondria elongate, are spared from degradation and sustain cell viability. Nat. Cell Biol. 13, 589–598.

15. Rambold, A.S., Kostelecky, B., Elia, N., and Lippincott-Schwartz, J. (2011). Tubular network formation protects mitochondria from autophagosomal degradation during nutrient starvation. Proc. Natl. Acad. Sci. U. S. A. 108, 10190–10195.

16. Rambold, A.S., Cohen, S., and Lippincott-Schwartz, J. (2015). Fatty acid trafficking in starved cells: regulation by lipid droplet lipolysis, autophagy, and mitochondrial fusion dynamics. Dev. Cell 32, 678–692.

17. Weir, H.J., Yao, P., Huynh, F.K., Escoubas, C.C., Goncalves, R.L., Burkewitz, K., Laboy, R., Hirschey, M.D., and Mair, W.B. (2017). Dietary Restriction and AMPK Increase Lifespan via Mitochondrial Network and Peroxisome Remodeling. Cell Metab. 26, 884–896.e5.

18. Villalobos, T.V., Ghosh, B., DeLeo, K.R., Alam, S., Ricaurte-Perez, C., Wang, A., Mercola, B.M., Butsch, T.J., Ramos, C.D., Das, S., et al. (2023). Tubular lysosome induction couples animal starvation to healthy aging. Nat Aging 3, 1091–1106.

19. Longo, V.D., and Panda, S. (2016). Fasting, circadian rhythms, and time-restricted feeding in healthy lifespan. Cell Metab. 23, 1048–1059.

20. Simpson, S.J., Le Couteur, D.G., Raubenheimer, D., Solon-Biet, S.M., Cooney, G.J., Cogger, V.C., and Fontana, L. (2017). Dietary protein, aging and nutritional geometry. Ageing Res. Rev. 39, 78–86.

21. Green, C.L., and Lamming, D.W. (2019). Regulation of metabolic health by essential dietary amino acids. Mech. Ageing Dev. 177, 186–200.

22. Bruss, M.D., Khambatta, C.F., Ruby, M.A., Aggarwal, I., and Hellerstein, M.K. (2010). Calorie restriction increases fatty acid synthesis and whole body fat oxidation rates. Am. J. Physiol. Endocrinol. Metab. 298, E108–16.

23. Aon, M.A., Bernier, M., Mitchell, S.J., Di Germanio, C., Mattison, J.A., Ehrlich, M.R., Colman, R.J., Anderson, R.M., and de Cabo, R. (2020). Untangling Determinants of Enhanced Health and Lifespan through a Multi-omics Approach in Mice. Cell Metab. 32, 100–116.e4.

24. Derous, D., Mitchell, S.E., Wang, L., Green, C.L., Wang, Y., Chen, L., Han, J.-D.J., Promislow, D.E.L., Lusseau, D., Douglas, A., et al. (2017). The effects of graded levels of calorie restriction: XI. Evaluation of the main hypotheses underpinning the life extension effects of CR using the hepatic transcriptome. Aging 9, 1770–1824.

25. Hansen, M., Rubinsztein, D.C., and Walker, D.W. (2018). Autophagy as a promoter of longevity: insights from model organisms. Nat. Rev. Mol. Cell Biol. 19, 579–593.

26. Aman, Y., Schmauck-Medina, T., Hansen, M., Morimoto, R.I., Simon, A.K., Bjedov, I., Palikaras, K., Simonsen, A., Johansen, T., Tavernarakis, N., et al. (2021). Autophagy in healthy aging and disease. Nat Aging 1, 634–650.

27. Tavernarakis, N., and Driscoll, M. (2002). Caloric restriction and lifespan: a role for protein turnover? Mech. Ageing Dev. 123, 215–229.

28. Madeo, F., Carmona-Gutierrez, D., Hofer, S.J., and Kroemer, G. (2019). Caloric restriction mimetics against age-associated disease: Targets, mechanisms, and therapeutic potential. Cell Metab. 29, 592–610.

29. Price, J.C., Khambatta, C.F., Li, K.W., Bruss, M.D., Shankaran, M., Dalidd, M., Floreani, N.A., Roberts, L.S., Turner, S.M., Holmes, W.E., et al. (2012). The effect of long term calorie restriction on in vivo hepatic proteostatis: a novel combination of dynamic and quantitative proteomics. Mol. Cell. Proteomics 11, 1801–1814.

30. Miller, B.F., Robinson, M.M., Reuland, D.J., Drake, J.C., Peelor, F.F., 3rd, Bruss, M.D., Hellerstein, M.K., and Hamilton, K.L. (2013). Calorie restriction does not increase short-term or long-term protein synthesis. J. Gerontol. A Biol. Sci. Med. Sci. 68, 530–538.

31. Dai, D.-F., Karunadharma, P.P., Chiao, Y.A., Basisty, N., Crispin, D., Hsieh, E.J., Chen, T., Gu, H., Djukovic, D., Raftery, D., et al. (2014). Altered proteome turnover and remodeling by short-term caloric restriction or rapamycin rejuvenate the aging heart. Aging Cell 13, 529–539.

32. Steinhauser, M.L., and Lechene, C.P. (2013). Quantitative imaging of subcellular metabolism with stable isotopes and multi-isotope imaging mass spectrometry. Semin. Cell Dev. Biol. 24, 661–667.

33. Ching, T.-T., and Hsu, A.-L. (2011). Solid plate-based dietary restriction in Caenorhabditis elegans. J. Vis. Exp. 10.3791/2701.

34. Froehlich, J.J., Rajewsky, N., and Ewald, C.Y. (2021). Estimation of C. elegans cell- and tissue volumes. MicroPubl. Biol. 2021, 10.17912/micropub.biology.000345.

35. Hall, D.H., Hartwieg, E., and Nguyen, K.C.Q. (2012). Modern electron microscopy methods for C. elegans. Methods Cell Biol. 107, 93–149.

36. Bélanger, S., Berensmann, H., Baena, V., Duncan, K., Meyers, B.C., Narayan, K., and Czymmek, K.J. (2022). A versatile enhanced freeze-substitution protocol for volume electron microscopy. Front Cell Dev Biol 10, 933376.

37. Lechene, C., Hillion, F., McMahon, G., Benson, D., Kleinfeld, A.M., Kampf, J.P., Distel, D., Luyten, Y., Bonventre, J., Hentschel, D., et al. (2006). High-resolution quantitative imaging of mammalian and bacterial cells using stable isotope mass spectrometry. J. Biol. 5, 20.

38. Krishna, S., Arrojo E Drigo, R., Capitanio, J.S., Ramachandra, R., Ellisman, M., and Hetzer, M.W. (2021). Identification of long-lived proteins in the mitochondria reveals increased stability of the electron transport chain. Dev. Cell 56, 2952–2965.e9.

39. Toyama, B.H., Arrojo E Drigo, R., Lev-Ram, V., Ramachandra, R., Deerinck, T.J., Lechene, C., Ellisman, M.H., and Hetzer, M.W. (2019). Visualization of long-lived proteins reveals age mosaicism within nuclei of postmitotic cells. J. Cell Biol. 218, 433–444.

40. Gurkar, A.U., Okawa, S., Guillermier, C., Chaddha, K., and Steinhauser, M.L. (2025). Large-scale clustered transcriptional silencing associated with cellular senescence. Aging Cell 24, e70015.

41. Min, K.-J., Hogan, M.F., Tatar, M., and O’Brien, D.M. (2006). Resource allocation to reproduction and soma in Drosophila: a stable isotope analysis of carbon from dietary sugar. J. Insect Physiol. 52, 763–770.

42. Hsin, H., and Kenyon, C. (1999). Signals from the reproductive system regulate the lifespan of C. elegans. Nature 399, 362–366.

43. Arantes-Oliveira, N., Apfeld, J., Dillin, A., and Kenyon, C. (2002). Regulation of life-span by germ-line stem cells in Caenorhabditis elegans. Science 295, 502–505.

44. Coscia, S.M., Moore, A.S., Thompson, C.P., Tirrito, C.F., Ostap, E.M., and Holzbaur, E.L.F. (2024). An interphase actin wave promotes mitochondrial content mixing and organelle homeostasis. Nat. Commun. 15, 3793.

45. Gugel, J., Currie, J., Alamillo, L., Flint, J., Kim, K.-Y., Debliqui, M., Ellisman, M.H., Lam, M.P.Y., Lau, E., Arrojo E Drigo, R., et al. (2026). Longevity of cardiac and skeletal muscle proteins is dependent on tissue and subcellular compartmentation patterns. Cell Rep. 45, 116768.

46. Dorn, G.W., 2nd (2019). Evolving concepts of mitochondrial dynamics. Annu. Rev. Physiol. 81, 1–17.

47. Morrow, C.S., Yao, P., Vergani-Junior, C.A., Anekal, P.V., Montero Llopis, P., Miller, J.W., Benayoun, B.A., and Mair, W.B. (2024). Endogenous mitochondrial NAD(P)H fluorescence can predict lifespan. Commun. Biol. 7, 1551.

48. Kuchel, G.A., Hevener, A.L., Ruby, J.G., Sebastiani, P., and Kumar, V. (2025). Workshop report-heterogeneity and Successful Aging Part I: Heterogeneity in aging-challenges and opportunities. J. Gerontol. A Biol. Sci. Med. Sci. 80, glaf023.

49. Salimi, S., Raftery, D., and Ferrucci, L. (2026). Genomic perspective on heterogeneity of organs and body aging. Aging Cell 25, e70353.

50. Palla, G., Spitzer, H., Klein, M., Fischer, D., Schaar, A.C., Kuemmerle, L.B., Rybakov, S., Ibarra, I.L., Holmberg, O., Virshup, I., et al. (2022). Squidpy: a scalable framework for spatial omics analysis. Nat. Methods 19, 171–178.

